# KHz-rate volumetric voltage imaging of the whole zebrafish heart

**DOI:** 10.1101/2020.07.13.196063

**Authors:** L. Sacconi, L. Silvestri, E.C. Rodríguez, G.A.B. Armstrong, F.S. Pavone, A. Shrier, G. Bub

## Abstract

Fast volumetric imaging is essential for understanding the function of excitable tissues such as those found in the brain and heart. Measuring cardiac voltage transients in tissue volumes with spatial and temporal resolutions needed to give insight to cardiac function has so far been impossible. We introduce a new imaging modality based on simultaneous illumination of multiple planes in the tissue and parallel detection with multiple cameras, avoiding compromises inherent in any scanning approach. The system enables imaging of voltage transients in-situ, allowing us, for the first time, to map voltage activity in the whole heart volume at KHz rates. The unprecedented spatio-temporal resolution of our method enabled the observation of novel dynamics of electrical propagation through the zebrafish atrioventricular canal.

---

Organs with excitable cells such as the heart and brain propagate voltage signals at high speed through spatially complex structures in three dimensions. Imaging activity in deep tissue volumes pose challenges that have been partially met in neural tissues by leveraging either the relative sparsity of active cells^1,2^ or rely on measuring slowly varying ions such as calcium^3^. These strategies are less viable in the heart, where cardiac function sensitively depends on waves of electrical activity rapidly propagating through dense layers of connected cardiac cells. Researchers have made headway measuring these waves using techniques that estimate activity in deep tissue using tomographic reconstruction^4,5^, or measure mechanical contraction as a surrogate for voltage using ultrasound^6^. The most promising approach for directly measuring voltage at cellular scales in tissue volumes is light-sheet microscopy (LSM) as it can capture confocal sections with the long integration times needed to measure activity from voltage sensitive dyes. LSM-based volumetric imaging strategies either rely on rapidly scanning the sheet through the sample^7–12^, use gating to capture information at different points in the cardiac cycle over many successive contractions^13,14^, or use parallel illumination planes to increase acquisition rates^15,16^. However, these approaches have fundamental limitations. Fast scanning methods trade integration time for high acquisition speed^8,11^, and gating methods can only image regularly repeating events^13^. Finally, current parallel plane strategies either greatly restrict the number and location^16^ of imaging planes, or, when multiplexing data from several planes onto a single detector^15^, must increase acquisition time proportionately to the number of planes to maintain signal quality. While some parallel plane architectures could, in principle,^17,18^ operate at high rates using specialized high speed cameras, so far there has been no demonstration of volumetric data capture with the signal quality needed to track voltage transients at physiologically relevant timescales.

Simultaneous Parallel Excitation and Emission Detection (SPEED) microscopy is a new technique that captures data from multiple light-sheets on multiple sensors in real time. Conventional light sheet microscopy achieves optical sectioning by ensuring that all fluorescence originates from a single plane. In our system, multiple parallel light sheets are projected into the sample and each plane is imaged by its own camera (Fig 1A). Each camera captures in-focus light originating from a single light sheet along with out-of-focus light from adjacent ones. Standard deconvolution algorithms are used to redirect out-of-focus signal to the correct plane to further improve optical sectioning. There are two key benefits of this configuration over other parallel-plane approaches. First, we can have many illumination planes that fully cover the imaging volume, allowing capture of z-stacks without scanning. Second, the use of multiple detectors allows for high acquisition rates without a specialized high-speed sensor. In addition, the multi-sensor architecture allows visualization of optical sections in real-time during experimental runs while avoiding trade-offs in signal quality, spatial resolution and dynamic range that are often associated with encoding data from multiple sources on one camera.

**Fig 1.**
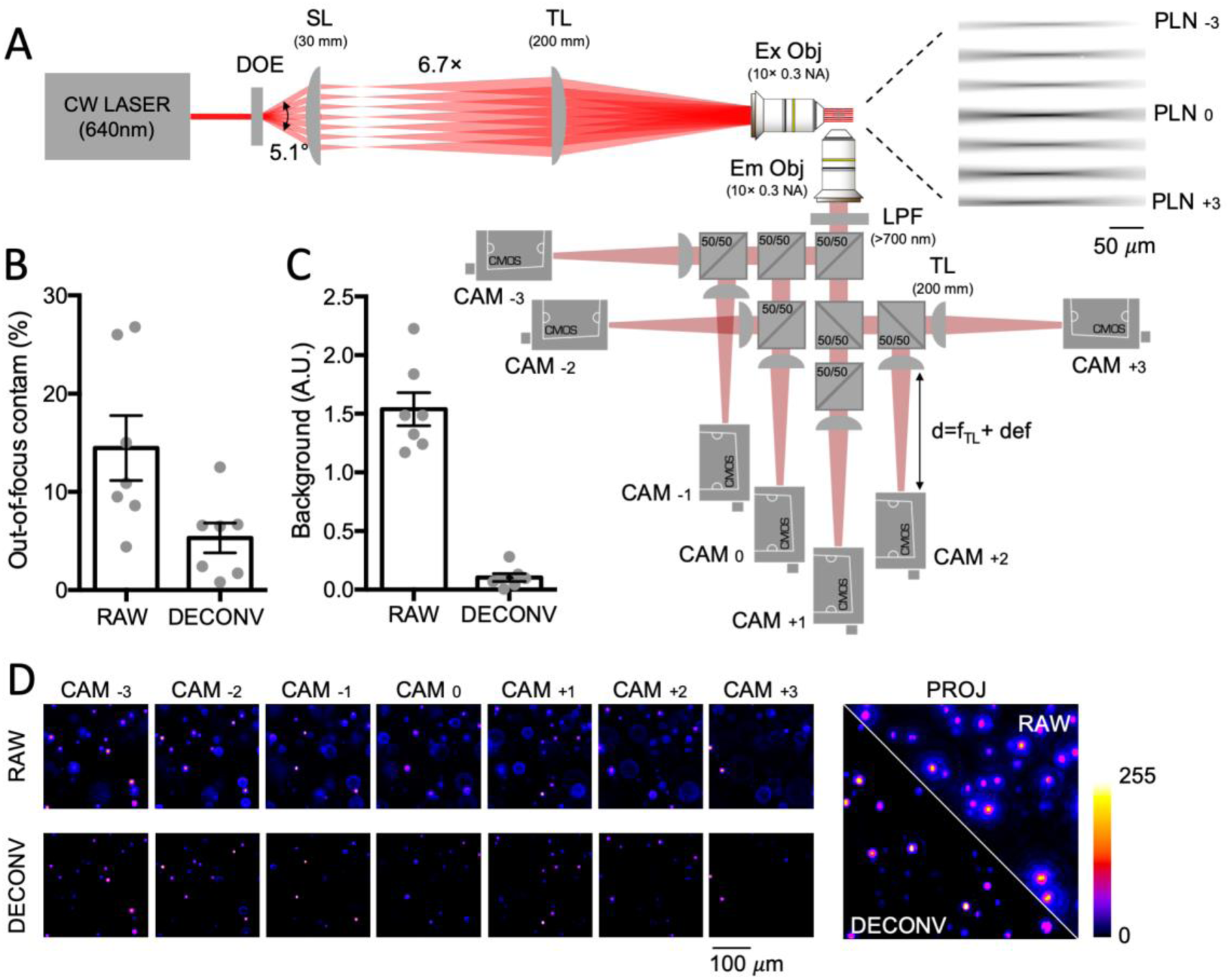
SPEED microscopy. A) Optical scheme: A 640 nm continuous wave (CW) laser beam is split and shaped as a light sheet by diffractive optical element (DOE) and coupled by scan lens (SL) and tube lens (TL) to a 10 X objective; the inset shows the real excitation light sheets imaged from an axis orthogonal to both excitation and detection, using a fluorescent gel; an orthogonally placed objective collects emitted fluorescence, which is then split by a cascade of 50:50 unpolarized beam splitters and focused by detection tube lenses (TL) onto cameras placed at different focal positions (f_TL_ + variable defocus); a long-pass filter (LPF) placed just after the detection objective blocks excitation light. A picture of the assembled system is shown in Supplementary Figure S2. B) Quantification of out-of-focus light contamination to images (raw vs deconvoluted); C) Quantification of background light intensity (raw vs deconvoluted); D) Images of fluorescent beads from the seven cameras (raw vs deconvoluted) with a zoomed view of a maximum-intensity projection.

We demonstrate SPEED using seven light sheets statically generated from a 645 nm laser light (OxxiusLBX-638-180-CSB-PPA) by a diffractive optical element (HoloeyeDE-R 251) and projected onto a (250×250×250)μm volume by a 10× objective (Fig 1A). Fluorescent images are collected by seven machine vision cameras (Basler acA720-520 μm) in parallel and processed using standard deconvolution algorithms (Huygens Professional), transferring signal to the correct image plane to further increase contrast and optical sectioning. We demonstrate the effectiveness of this approach using a sample phantom containing fluorescent beads (Fig 1B-D).

The zebrafish is emerging as a popular experimental model for cardiac research, in part because the heart has voltage transients that are similar to those in human and its small size makes the entire heart accessible to deep tissue imaging techniques that allow in-situ imaging of embedded structures. Current deep tissue imaging modalities are, however, too slow to capture electrical activity at millisecond resolution with sufficient spatial detail to image propagation in the zebrafish heart. We positioned explanted zebrafish hearts loaded with red-shifted voltage sensitive dye (Di-4-ANBDQPQ,1 μM) into the SPEED microscope as shown in Fig 2A. Fluorescence was captured from seven 400×400 pixel planes (Fig 2A,D) simultaneously at over 500 frames/s, allowing us to map activity in a (250 μm)^3^ volume (Fig 2B,C). Action potentials from three regions of interests (ROI) are shown for the raw and deconvolved data sets in Fig 2E, with one ROI positioned over atrial tissue (blue), one over the ventricular tissue (red), and a third positioned in tissue between these (green) in a region that corresponds to the atrioventricular (AV) canal. While action potential morphology for atrial (blue) and ventricular (red) ROIs is clearly resolvable in the raw dataset, the AV action potential shape is corrupted by signal from overlapping layers.

**Fig 2.**
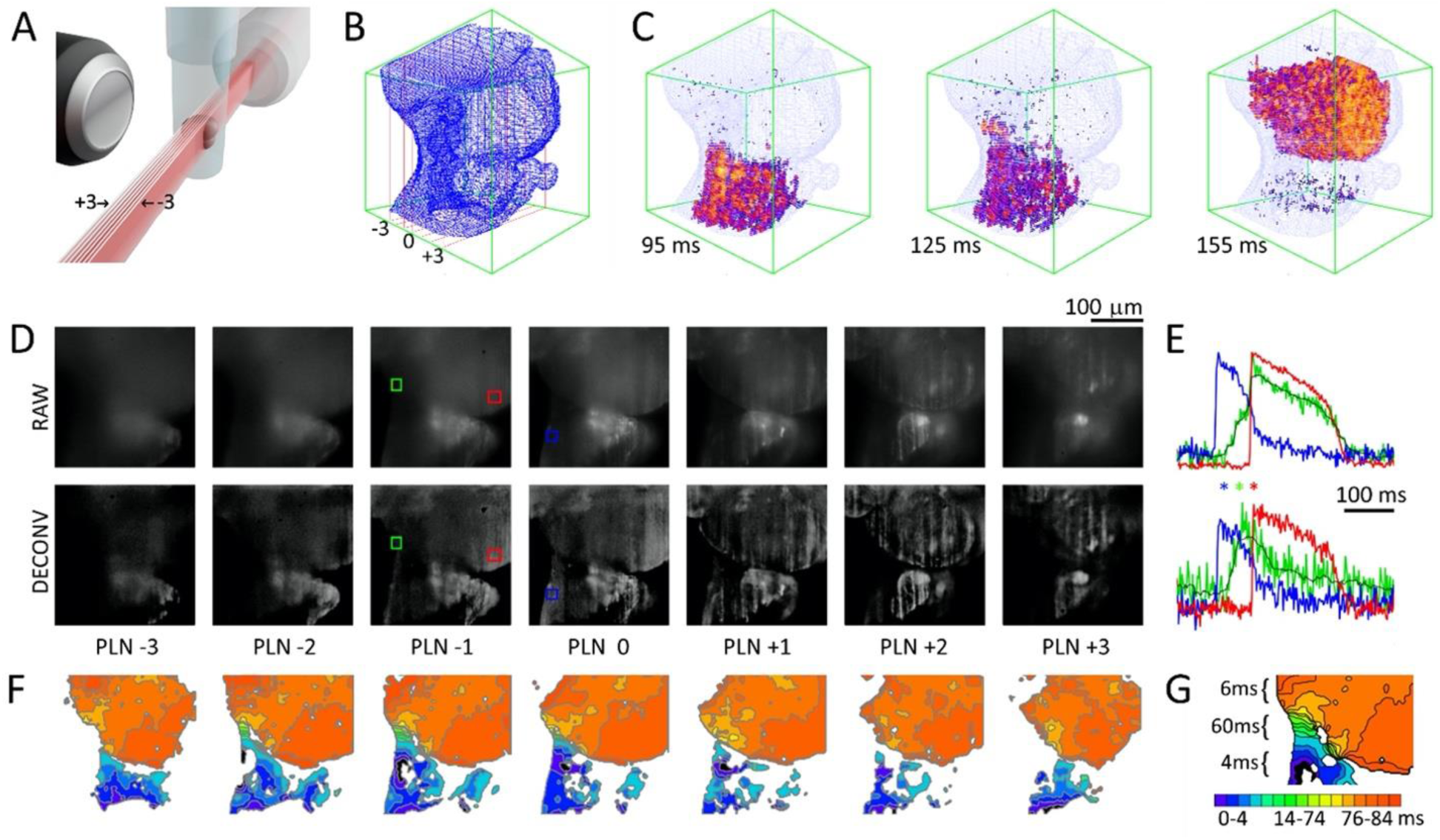
A) Orientation of sample relative to the seven light sheets; B) Surface mesh generated from high resolution z-stack; C) Propagation of voltage transients in the seven planes at 95ms, 125ms and 155ms; D) Comparison of the seven planes (PLN-3 through PLN+3) for raw and deconvoluted data sets; with ROIs in PLN −1 and 0; E) Action potential morphology from atrium (blue), ventricle (red) and AV canal region (red) for raw and deconvolved datasets from the ROIs in D. Blue, green and red asterisks symbols correspond to 95ms, 125ms and 155ms to show the relative timing in the maps shown in panel ‘C’; F) Isochronal maps show propagation through atrial, AV, and ventricular tissues; G) Inset shows a map of the slow conduction through the AV region.

Deconvolution reveals a voltage transient morphology in slowly conducting tissue that is distinct from that found in atrial or ventricular tissues. The deconvolved data set is then used to generate isochronal maps showing propagation in each plane, with detail sufficient to map conduction through the slowly conducting AV canal (Fig 2F,G). Propagation in the entire imaged volume is then visualized in 3D by overlaying the seven planes on a conventionally acquired z-stack (Fig 2B, C and Supplementary movie S1).

SPEED’s light efficiency allows long record times without apparent phototoxicity or bleaching, enabling data capture for over ninety minutes with little loss in signal quality. We leveraged the system’s ability to capture long records to make an experimental model of an arrhythmia where waves backpropagate from the ventricles to the atria. Early afterdepolarizations (EADs) occur in myocytes due to imbalances in cardiac action potential repolarization kinetics. In certain circumstances EADs can trigger a macroscopic propagating wave that results in a premature contraction; EADs are associated with potentially fatal arrhythmias and are thus active areas of research. Repolarization kinetics were pharmacologically altered in the zebrafish hearts (online methods) resulting in prolonged ventricular action potentials (Fig. 3D) and EADs (Fig 3E). EADs generate extrasystolic waves that back-propagate through the AV canal and re-excite the atrium (Fig 3B,C,G and Fig S2). Conduction times between the ventricle and atrium are consistent but slightly longer (Fig 3F,G) than those measured between the atria and ventricles during normal rhythms, suggesting that the AV canal is optimized for propagating waves in one direction or that the AV tissue is only partially repolarized. Interestingly, increased ventricular to atrial conduction times are also observed in human hearts^19^. Further study is needed to determine whether the mechanisms for direction dependent conduction times in these systems are the same.

**Fig 3.**
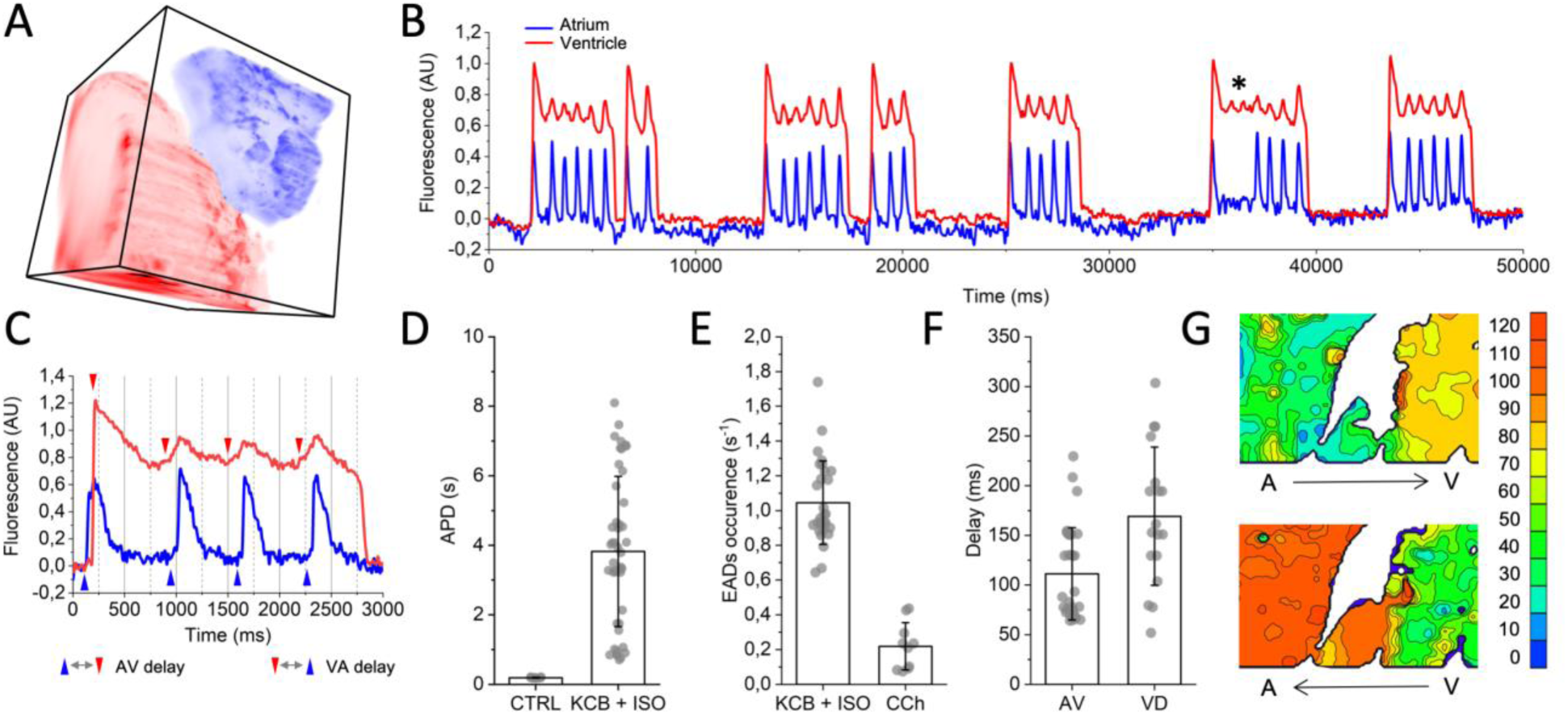
A) volume map of atrial (blue) and ventricular (red) regions obtained from high resolution z-stack. B) Action potentials from ventricular and atrial regions show backpropagation of ventricular early afterdepolarizations (EAD) activating atrial tissue (star: EADs without backpropagation). C) Ventricular EAD onset (red arrows) occur before atrial activations (blue arrows); D) Effect of E4031 (KCB) and Isoproterenol (ISO) on action potential duration in the ventricle; E) EADs per action potential over all records in E4031 and isoproterenol treated hearts – EAD frequency greatly reduced by carbachol (CCh); F) delay between atrium and ventricle is slightly lower than the backpropagation delay between the ventricle and atrium. G) Isochronal maps of a 170 × 100 μm cardiac section with isochrones drawn every 10ms show atrial (A) to ventricular (V) propagation direction for the first atrial beat in Fig C (top) and back-propagation from the ventricle to the atria for subsequent EAD triggered beats (bottom). See Supplementary Figure S4 for the entire 250 × 250 μm field of view.

SPEED’s low phototoxicity and high signal quality is a consequence of having an architecture that maximizes sensor integration times using true parallel volumetric acquisition. Light-sheet imaging has advantages in capturing dynamic events from weakly fluorescent tissue as exposure times for each pixel can be far longer than scanning system at a similar resolution^20^. SPEED’s architecture extends light-sheet’s simultaneous imaging paradigm from 2D to 3D, allowing capture of z-stacks in thick samples in a single camera exposure. The use of multiple cameras removes frame-rate bottlenecks associated with using a single large sensor while allowing visualization of optical sections during acquisition, which can be improved offline with deconvolution.

We demonstrated our imaging system in zebrafish hearts as its popularity as model system for studying cardiac dynamics is growing. Indeed, there are several studies that use voltage sensitive dyes to map propagation in the zebrafish heart^21–24^, but all rely on measuring activity at the surface, and none are able to image interior structures such as the AV canal. SPEED can both capture fast dynamics of deep structures (Fig 2) and perform long-duration experiments (Fig 3) in cardiac preparations. While we have optimized SPEED for imaging zebrafish cardiac dynamics, it can be used for high speed volumetric imaging of any excitable tissue including the central nervous system. With suitable optimization, SPEED could open the possibility to directly investigate electrical activity across neuronal populations, elucidating computational rules that remain hidden in the low temporal resolution of calcium indicators^25,26^. Although the system is not restricted to a specific illumination wavelength, our implementation with red light paves the way for effective coupling with optogenetic stimulation with blue light with negligible cross-talk. The unique possibility of combining fast volumetric voltage imaging with the genetic specificity of channelrhodopsin opens research possibilities that are difficult to overestimate. For instance, spatio-temporal neuronal activation patterns related to a specific stimulus or experience could be reproduced using three-dimensional holographic photostimulation^27^ to investigate complex neuronal networks using a truly electrical pump-and-probe approach. A similar approach could be applied to cardiovascular research to disentangle the various roles that nonmyocyte cardiac cell populations^28–30^ have in modulating cardiac wave dynamics in physiological and pathological conditions.

## Materials and Methods

### Zebrafish

Wild-type AB zebrafish (*Danio rerio*) were bred and maintained according to standard procedures^31^. All experiments were performed in compliance with the guidelines of the Canadian Council for Animal Care and conducted at McGill University. Experiments were performed on sexually undifferentiated zebrafish larvae aged 2 weeks post-fertilization. Hearts were extracted from larvae and kept at room temperature in standard fish system water with 10 mM glucose.

### Dye loading and imaging

Excised hearts were placed in 1 μM Di-4-ANBDQPQ in Evans for 5 minutes and were placed in a dish containing 2% solution low melting point agarose (Sigma) dissolved in Evans solution. As the gel started to set, the heart was positioned into a glass capillary tube. A ∼1mm segment of agar containing the heart was pushed out and suspended from the capillary tube in a square walled plastic cuvette containing 1ml Evans for recording. For most experiments, 5 μM blebbistatin is added to the agarose and recording solution to arrest contraction. For the experiments presented in Figure 3, potassium channel blocker E4031 (final concentration 20 μM), cholinergic agonist carbachol (final concentration 1 μM) and β-adrenoreceptor agonist isoproterenol (final concentration 1 μM) are added directly to the cuvette from diluted stock solutions. Isoproterenol induced afterdepolarizations by activating the cardiac beta receptor, leading to an increase in cAMP via upregulation of adenylyl cyclase activity. Isoproterenol’s role was confirmed by blocking its effect using carbachol, which activates muscarinic receptors resulting in downregulation of adenylyl cyclase.

### Software - hardware control

The seven cameras were controlled using custom written software interface written in java and Basler’s free Pylon SDK. The software allows simultaneous viewing of the seven cameras for focusing and acquisition. Alignment was achieved by writing a wrapper around Fiji plugin (Register_Virtual_Stack_MT.java; https://imagej.net/Register_Virtual_Stack_Slices) to find pixel offsets for each camera after focusing on a high contrast image target, which were then used to select a region from each camera for recording. The number of pixels in the recording region sets bounds on the frame rate due to data transfer limitations in the USB cameras used in this study: we use a 400×400 pixel region for 500 Hz imaging a 128×128 pixel region and image at 1000Hz (see Figure S3). During image acquisition, cameras were synchronized to a timing pulse set using an Arduino Uno microcontroller, which also triggered LED and laser illumination. A set of 1000 frames from each camera was saved in a circular buffer and transferred to disk as a single file for each record.

### Analysis

Data was deconvolved by first converting image data to tiffs and processing using commercial software (Huygens Professional version 19.04, Scientific Volume Imaging; Algorithm used: Classical Maximum Likelihood Estimation, 40 iterations with a theoretical PSF). 3D meshes (Fig 3B) were generated using Fiji’s 3D viewer plugin (v 4.02, https://imagej.net/3D_Viewer) and exported into custom written software that extended Processing’s (https://www.processing.org) PApplet class and its P3D openGL environment.

Isochronal maps were generated by spatially binning data to give 200×200 pixel images for each view, normalizing data so that temporal transients for each pixel have the same magnitude, filtering the data with a running five frame temporal and five pixel radius spatial median filters prior to using a marching squares algorithm to locate wavefronts based on the threshold-crossing time for each pixel. For figure 3G, S2 and movie S3, backpropagation was measured by averaging the signals from two cameras and averaging signals from three successive EADs prior to generating the isochronal maps.

## Acknowledgements

G. Bub acknowledges support from the Canadian Heart and Stroke Foundation and the Canadian Foundation for Innovation. L. Sacconi acknowledges EMBO for supporting his visit to Bub’s laboratory with a short-term fellowship. The authors are grateful to Alex Quinn, Matthew Stoyek, Leon Glass, Michael Guevara, Claire Brown, Arjun Krishnaswami and Giuseppe Sancataldo for scientific advice and technical support throughout the project.

## Author Contributions

L. Sacconi: original concept; G. Bub and L. Sacconi: system construction and programming; G.A.B. Armstrong and E.C.Rodríguez: cardiac sample preparation; G. Bub, L. Sacconi and A. Shrier: designed and performed experiments; G. Bub, L. Sacconi and L. Silvestri: data analysis; F.S. Pavone contributed new analytic tools; all authors contributed to writing the document and data interpretation.

