## Supplemental figures for "KHz-rate volumetric voltage imaging of the whole zebrafish heart"

Supplementary figures

Supplemental Movie S1: Video showing propagation through the atria, AV canal region and ventricles, imaged at 500 Hz. The volume imaged corresponds to a 250x250x250 μm cube of the spontaneously beating zebrafish heart.

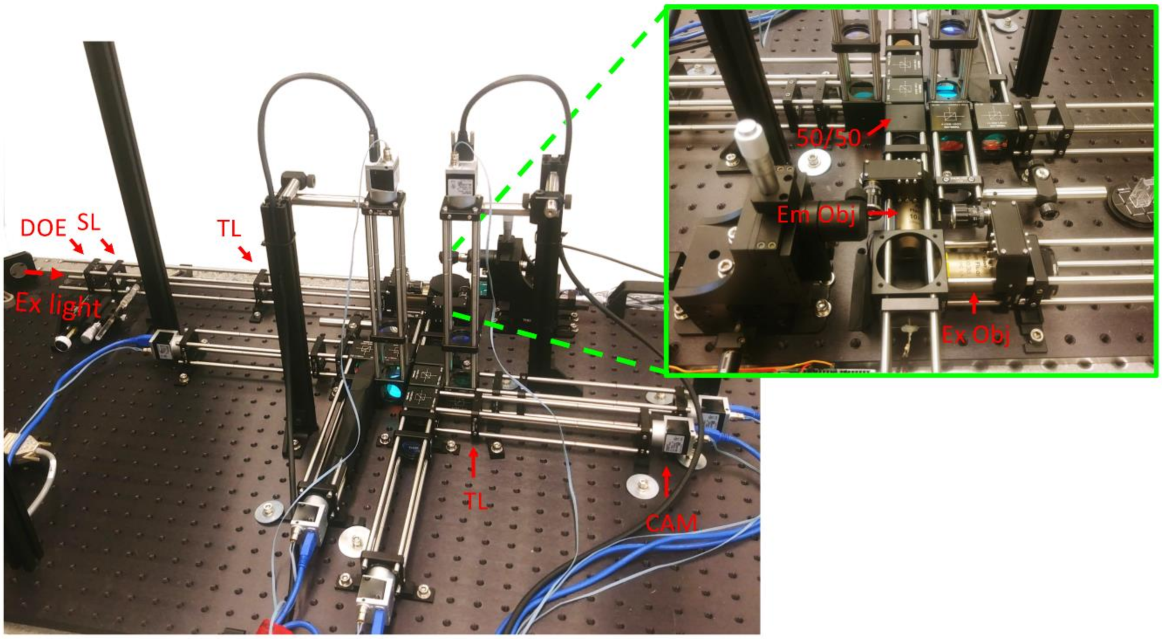


**Supplemental Figure S2**: Image of the SPEED microscope, showing the seven cameras (CAM), tube lenses (TL), diffractive optical element (DOE), scan lens (SL) and light path. The system is constructed using standard Thorlabs beam splitting cubes and 30mm cage components. The inset shows the imaging area with orthogonally placed objectives and a series of 50/50 beam splitting cubes. See Figure 1 (main text) for details.

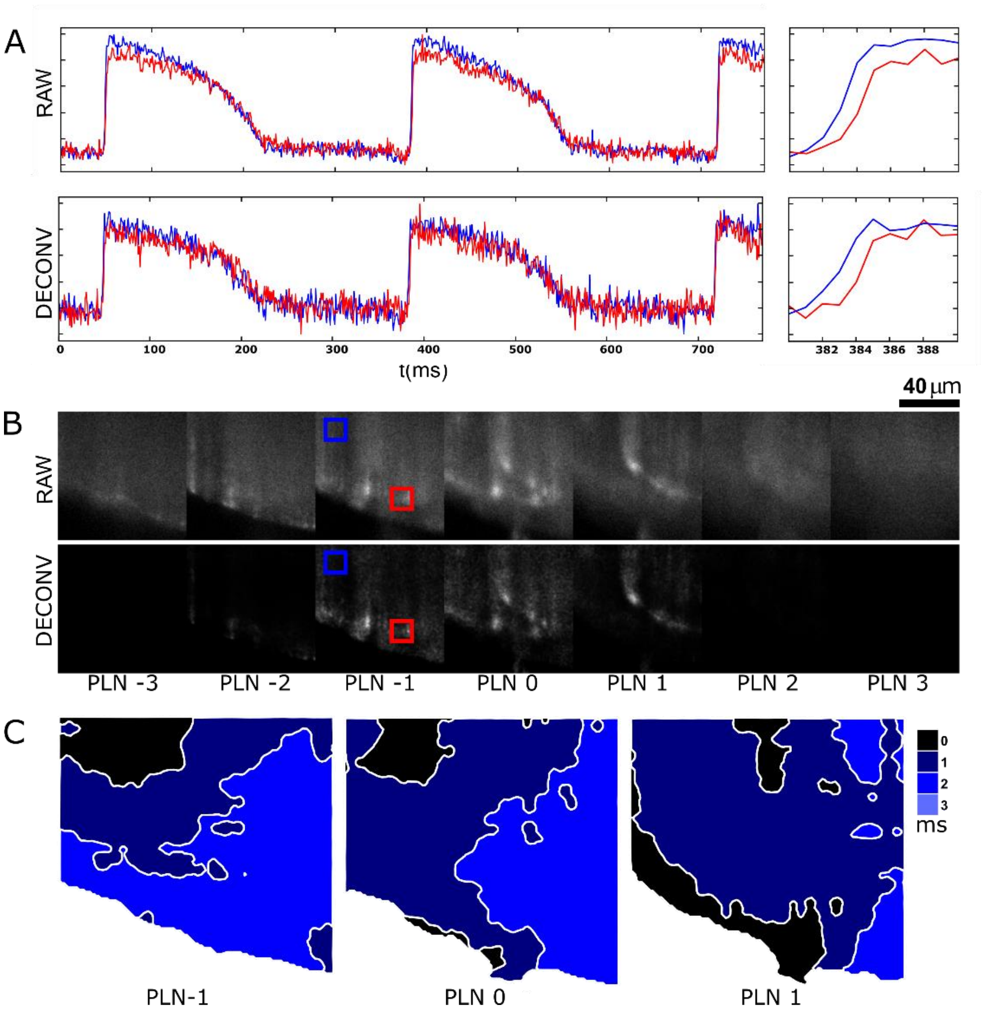


**Supplemental Figure S3**: 1000 f/s (KHz-rate) imaging of ventricular tissue. 128x128 pixel region was selected to image propagation across the ventricle at high speeds. Intensity vs time traces (panel A) show propagation across the field occurs over approximately 2ms, for both the raw and deconvolved signal. Deconvolution (panel B, bottom) resulted in a redistribution of signal to three out of the seven frames. Isochronal maps from the three central planes suggests that the wave propagates in both radially and axially.

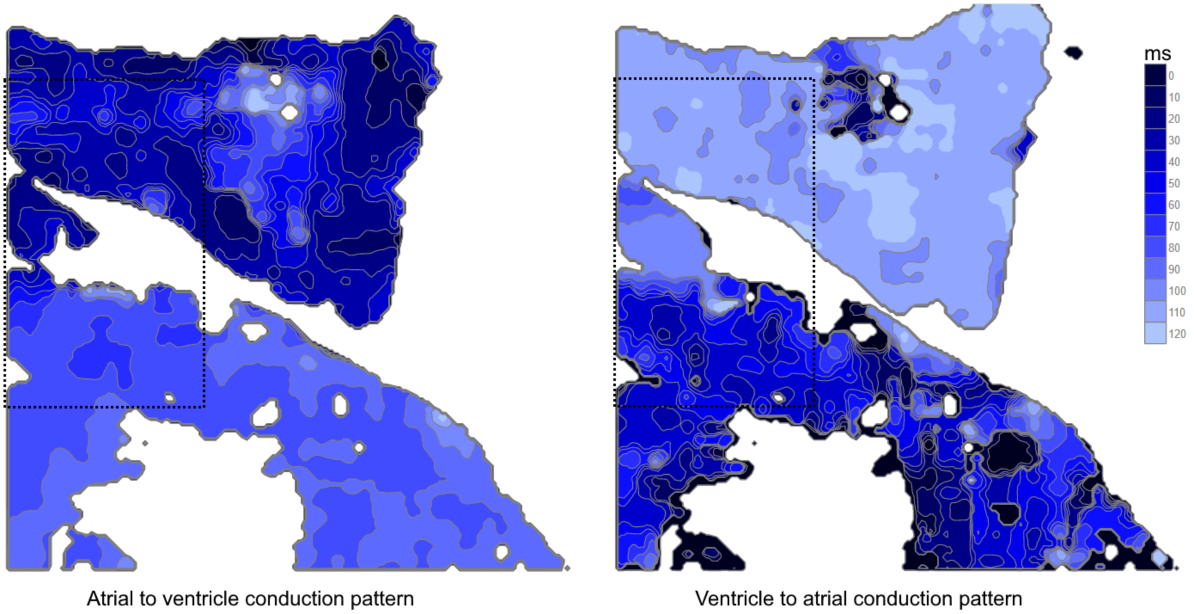


**Supplemental Figure S4**: Isochronal maps at two time points for a 250 x 250 μm region showing propagation during EAD dynamics for the first atrial beat, where the impulse originates in the atria and propagates via the AV canal to the ventricle (left panel), and for subsequent EAD beats, where the impulse originates in the ventricle and propagates retrogradely through the AV canal to activate the atria. The dashed line insets show the regions depicted in Figure 3g in the main text. Isochronal maps for EAD retrograde propagation were obtained by averaging signals from two adjacent cameras and averaging three successive EADs.
